# Methicillin resistant *Staphylococcus aureus* emerged long before the introduction of methicillin in to clinical practice

**DOI:** 10.1101/122408

**Authors:** Catriona P. Harkins, Bruno Pichon, Michel Doumith, Julian Parkhill, Henrik T. Westh, Alexander Tomasz, Herminia de Lencastre, Stephen D. Bentley, Angela M. Kearns, Matthew T.G. Holden

## Abstract

The spread of drug-resistant bacterial pathogens pose a major threat to global health. It is widely recognised that the widespread use of antibiotics has generated selective pressures that have driven the emergence of resistant strains. Methicillin-resistant *Staphylococcus aureus* (MRSA) was first observed in 1960, less than one year after the introduction of this second generation β-lactam antibiotic into clinical practice. Epidemiological evidence has always suggested that resistance arose around this period, when the *mecA* gene encoding methicillin resistance carried on an SCC*mec* element, was horizontally transferred to an intrinsically sensitive strain of *S. aureus*. Whole genome sequencing a collection of the very first MRSA isolates allowed us to reconstruct the evolutionary history of the archetypal MRSA. Bayesian phylogenetic reconstruction was applied to infer the time point at which this early MRSA lineage arose and when SCC*mec* was acquired. MRSA emerged in the mid 1940s, following the acquisition of an ancestral type I SCC*mec* element, some fourteen years prior to the first therapeutic use of methicillin. Methicillin use was not the original driving factor in the evolution of MRSA as previously thought. Rather it was the widespread use of first generation β-lactams such as penicillin in the years prior to the introduction of methicillin, which selected for *S. aureus* strains carrying the *mecA* determinant. Crucially this highlights how new drugs, introduced to circumvent known resistance mechanisms, can be rendered ineffective by unrecognised adaptations in the bacterial population due to the historic selective landscape created by the widespread use of other antibiotics.

## Introduction

Methicillin resistant *Staphylococcus aureus* (MRSA) has been identified as one of the major risk pathogens associated with the development of antimicrobial resistance (AMR). The emergence of AMR in *S. aureus* is well documented and the species has proven particularly adept at evolving resistance in the face of new antibiotic challenges. The introduction of penicillin in the 1940s heralded a revolution in the treatment of infectious diseases. However at the same time as its use was becoming more widespread following advances in the scaling up of production, evidence of penicillin resistance in *S. aureus* was already being uncovered (Kirby 1944).

Methicillin (Celbenin), a semi-synthetic β-lactam, was introduced in the UK in 1959 to circumvent growing penicillin resistance in *S. aureus*, associated with the acquisition of a β-lactamase enzyme, *blaZ* (Knox 1960). As a second-generation β-lactam antibiotic, methicillin was insensitive to breakdown by BlaZ. Following the introduction of methicillin into clinical practice in the UK the Staphylococcal Reference Laboratory in Colindale (London, England) screened *S. aureus* isolates for evidence of resistance to this antibiotic (Jevons 1961). More than 5000 *S. aureus* strains were assessed in the period between October 1959 and November 1960, and in October 1960 three isolates showing increased minimum inhibitory concentrations (MICs) to the new drug, methicillin, were identified. The isolates originated from the same hospital and shared a common phage type and resistance profile (penicillin, streptomycin and tetracycline) suggesting that they were related. In the description of these isolates it was noted that methicillin had been used only once previously at this hospital, and that none of the individuals from whom MRSA was isolated had been exposed to the drug. Within two years MRSA was being detected elsewhere in Europe, with invasive infections being identified in Denmark (Eriksen and Erichsen 1964). These MRSA isolates from the UK and Denmark in the early 1960s constitute the very first epidemic MRSA clone.

The genetic basis of methicillin resistance in *S. aureus* is associated with carriage of a mobile cassette of genes known as the Staphylococcal cassette chromosome *mec* (SCC*mec*) (Katayama et al. 2000). Within this cassette is the *mecA* gene that is responsible for resistance to β-lactams including methicillin. The product of *mecA* is the peptidoglycan synthesis enzyme penicillin binding protein (PBP) 2a involved in cross-linking of peptidoglycan in the bacterial cell wall (Hartman and Tomasz 1984; Matthews and Tomasz 1990). PBP2a has a lower binding affinity for β-lactam antibiotics than the native PBP proteins encoded in the core genome of *S. aureus*. The subsequent combination of reduced penicillin-binding affinity and increased production PBP2a accounts for the observed resistance to β-lactam antibiotics.

Genetic analyses of the first MRSA by multi-locus sequence typing (MLST) demonstrated that they were sequence type (ST) 250, a lineage belonging to clonal complex (CC) 8 and carried the type I SCC*mec* element (Crisóstomo et al. 2001; Enright et al. 2002). After emerging in the UK, this first epidemic MRSA clone (ST250-MRSA-I) spread across Europe during the 1960s and 70s, but by the late 1980s had become less prevalent and is now rarely reported (Enright et al. 2002; Oliveira et al. 2002; Gomes et al. 2006). The single locus variant and close relative of ST250-MRSA-I, ST247-MRSA-I was first detected in Denmark 1964 (Crisóstomo et al. 2001) and has been more successful, spreading globally and persisting as a source of outbreaks in Europe into the late 1990s (Oliveira et al. 2002; Gomes et al. 2006), but this too has been superseded by more successful contemporary clones (Oliveira et al. 2002). Five decades on since the appearance of the first MRSA, multiple MRSA lineages have emerged which have acquired different variants of SCC*mec* elements.

Epidemiological evidence has always suggested that MRSA arose following, and probably as a consequence of, the introduction of methicillin into clinical practice. However exactly when SCC*mec* was acquired by *S. aureus* has never before been determined. Here we have used whole genome sequencing of a collection of 209 of the earliest MRSA isolates recovered in Europe between 1960 and 1989 to reconstruct the evolutionary history of methicillin resistance. Using Bayesian phylogenetic reconstruction we have identified the likely time point at which this early lineage arose and also predicted the time by which SCC*mec* was acquired.

## Results

### Early MRSA belong to a diverse clone

Preserved in the culture collection of the Staphylococcal Reference Laboratory at Public Health England are representatives of the very first MRSA identified. These original isolates have been preserved as freeze-dried cultures, and have not been repeatedly passaged over the years. One hundred and eighty eight isolates that represented the earliest MRSA were recovered from the ampoules and their genomes sequenced (Supplementary Table 1). All the isolates belonged to CC8 MRSA and were originally isolated between 1960 and the late 1970s, and included eight isolates from the original study describing MRSA in 1961 (Jevons 1961). In addition, 21 CC8 MRSA isolated between 1964 and 1989 in Denmark (Crisóstomo et al. 2001; Gomes et al. 2006) were sequenced, as representatives of the earliest MRSA detected elsewhere in Europe. We also included early methicillin sensitive isolates of ST250 or ST247 (n=11), however only a limited number of these were found in the reference laboratory collection.

Analysis of the MLST of the isolates identified two main groups, ST250 (n=126), and a single locus variant (SLV), ST247 (n=78), plus two novel SLVs of ST247 (n=4) (Supplementary Table 1). A supplementary isolate from the Public Health England collection was included to provide an outgroup for the analysis;

RH12000692_7401696 is an MRSA which was collected in 1967 and is a triple locus variant of ST250 (Supplementary Table 1).

The *S. aureus* isolate COL, a representative member of this early MRSA lineage first identified in the 1960s (Dyke et al. 1966) had previously been fully sequenced, and the chromosome was used as a reference for mapping. Following exclusion of mobile genetic elements (MGEs) and predicted recombination events in the collection, a total of 4220 SNPs were identified and used to construct a phylogeny (Figure 1A). The population framework revealed a diverse population structure containing several distinct clades. The mapping of the ST information on to the phylogeny reveals that the ST250 population is basal to the ST247, suggesting that ST247 emerged from ST250, which is consistent with the epidemiological evidence, and supports the hypothesis that this pandemic multidrug-resistant MRSA clone emerged out of the ancestral MRSA genotype (Crisóstomo et al. 2001; Enright et al. 2002).

Highlighted in the expanded view (Figure 1B) are the isolates from the Jevons study, derived from three individuals at the same hospital in the South London area between July and November 1960 (Jevons 1961). The isolation source and resistance profiles of these isolates are shown in Supplementary Table 2. These isolates are genetically very closely related differing by seven SNPs only. Present within this cluster are additional isolates from the Public Health England collection originating between 1960 and 1961. Full epidemiological data is not available for these, but two of these isolates were identified in the same region as the hospital where the original Jevons study isolates originated. The genetic distance between isolates and their phylogenetic relationships suggests there was transmission within the hospital between patients A and C and nurse B, and that they were also transmitted beyond the hospital as part of a local outbreak.

Although all of the Jevons isolates are confined to a single clade, other isolates from the early 1960s are distributed throughout the entire phylogeny (Figure 2). This suggests that the earliest MRSA circulating in the UK were not from a single recently emerged clone, but were representatives of an established population. In addition to the UK isolates, there were 21 from Denmark, which represent the earliest MRSA detected outside the UK. These derive from 1964 onwards, and include the youngest isolates within the collection from the late 1980s. The Danish isolates are found in three clusters distributed throughout the phylogeny (Figure 1A) suggesting that, like the early UK MRSA, they originated from an established and diverse population.

**Figure 1.**
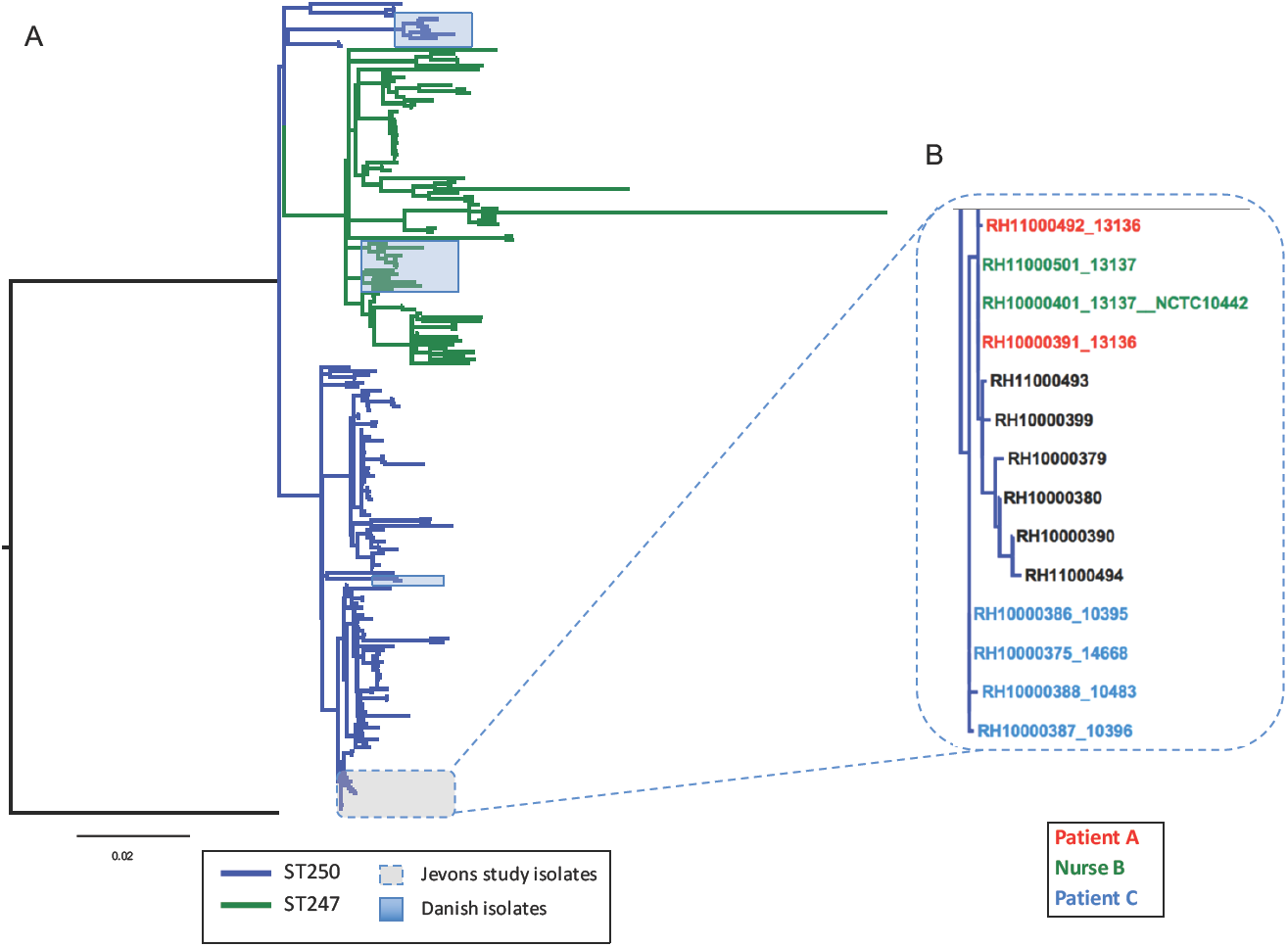
Population structure of historic MRSA isolates. A. Maximum likelihood tree of historic MRSA isolates. The tree was built using a maximum likelihood method using SNPs from the core genome of 209 isolates. Included in the phylogeny is the COL reference isolate to which the sequence reads were mapped. The tree is rooted with RH12000692_7401696 as an outgroup; this is a CC8 isolate and is a triple locus variant of ST250. Tree branches are coloured according to their ancestral sequence type population; blue indicate the ST250 and the green branches indicate the ST247. Isolates from Denmark are highlighted in the green shading and isolates described in the Jevons study are outlined in the dashed box, and a zoomed in view of the phylogeny is displayed in panel B. The coloured branch labels indicate the three individuals who supplied the original isolates in the Jevons study.

### Genetic basis of resistance to methicillin and other antibiotics in the archetypal MRSA population

Previous studies have shown that the archaic MRSA clone carried a type I SCC*mec* element, which was the first type of this MGE family to be classified (Katayama et al. 2000; Ito et al. 2001). Notably the description of the type I element was based upon the SCC*mec* derived from *S. aureus* strain NCTC10442 identified in the 1960 Jevons study (Supplementary Table 2 and Figure 1B) (Ito et al. 2001). The type I element carries *mecA* as its only resistance gene in combination with a truncated gene encoding the MecRI regulatory protein (together known as a class B *mec* gene complex) with type 1 chromosomal recombinases (*ccrA1* and *ccrB1*). The original description of SCC*mec* type I identified the presence of a frameshift mutation in *ccrB1* which disrupts the translation of this site specific recombinase (Ito et al. 2001). In the collection, 193 of the isolates contained intact SCC*mec* elements carrying the *mecA* gene (Figure 2). Of these 192 were SCC*mec* type I elements, all of which contained the NCTC10442-type frameshift mutation in *ccrB1*. The only non-type I element identified in the collection was in the outgroup isolate RH12000692_7401696, which contained a type IVh SCC*mec* element. The remaining 16 isolates lacking SCC*mec* elements were distributed throughout the phylogeny (Figure 2), suggesting that these represent MSSA arising from the loss of the type I SCC*mec* element, rather than forming an ancestral MSSA population.

In addition to methicillin resistance, the first MRSA described were also resistant to penicillin, streptomycin and tetracycline (Jevons 1961). Analysis of the genomes of these isolates identified *blaZ* and *tetK* genes conferring resistance to penicillin and tetracycline respectively, but failed to identify the *str* or *aad9* genes associated with streptomycin resistance in *S. aureus*. In the absence of an acquired resistance gene, the core genome was examined for mutations potentially responsible for resistance to streptomycin. In *Mycobacterium tuberculosis*, mutations in the ribosomal protein RpsL were shown to confer streptomycin resistance, including the substitution of an arginine for a lysine residue at residue 43 (Huang et al. 2003). Alignment of the *M. tuberculosis* and *S. aureus* sequences revealed that RpsL in the Jevons isolates contained an arginine in the equivalent position, residue 56. Comparison with RpsL sequences in the public sequence databases showed that in *S. aureus* the frequent amino acid residue at position 56 was lysine. Examining the whole collection, all but one of the sequenced isolates contained the arginine residue at position 56, the exception being the outgroup isolate RH12000692_7401696 (Figure 2). This demonstrates that the non-synonymous substitution resulting in an arginine for a lysine residue at residue 56 (K56R) occurred most likely very early during emergence of the archetypal MRSA population.

Analysis of the resistomes of the isolates revealed genetic resistance determinants to numerous other antibiotics, including: penicillin (*blaZ*), erythromycin (*ermA* and *linA*), kanamycin (*aadD*), gentamicin and kanamycin (*aacA*-*aphD*), spectinomycin and streptomycin (*aad9*), and chloramphenicol (*catA1, catA2* and *catA3*), fusidic acid (*fusA* P404L) and trimethoprim (*dfrA* F99Y), as well as well as genes associated with decreased susceptibility to disinfectants (*qacA* and *qacC*). The frequency and widespread dispersal of these determinants reveals the strong selective pressure exerted by antibiotics on the archetypal MRSA clone over an extensive period. Examining their distribution in the context of the phylogeny shows that some of these traits have been co-acquired (Figure 2), such as *ermA* and *aad9* which are carried on Tn*554*, and that these acquisition events can be mapped on to the phylogeny (Murphy et al. 1985).

**Figure 2.**
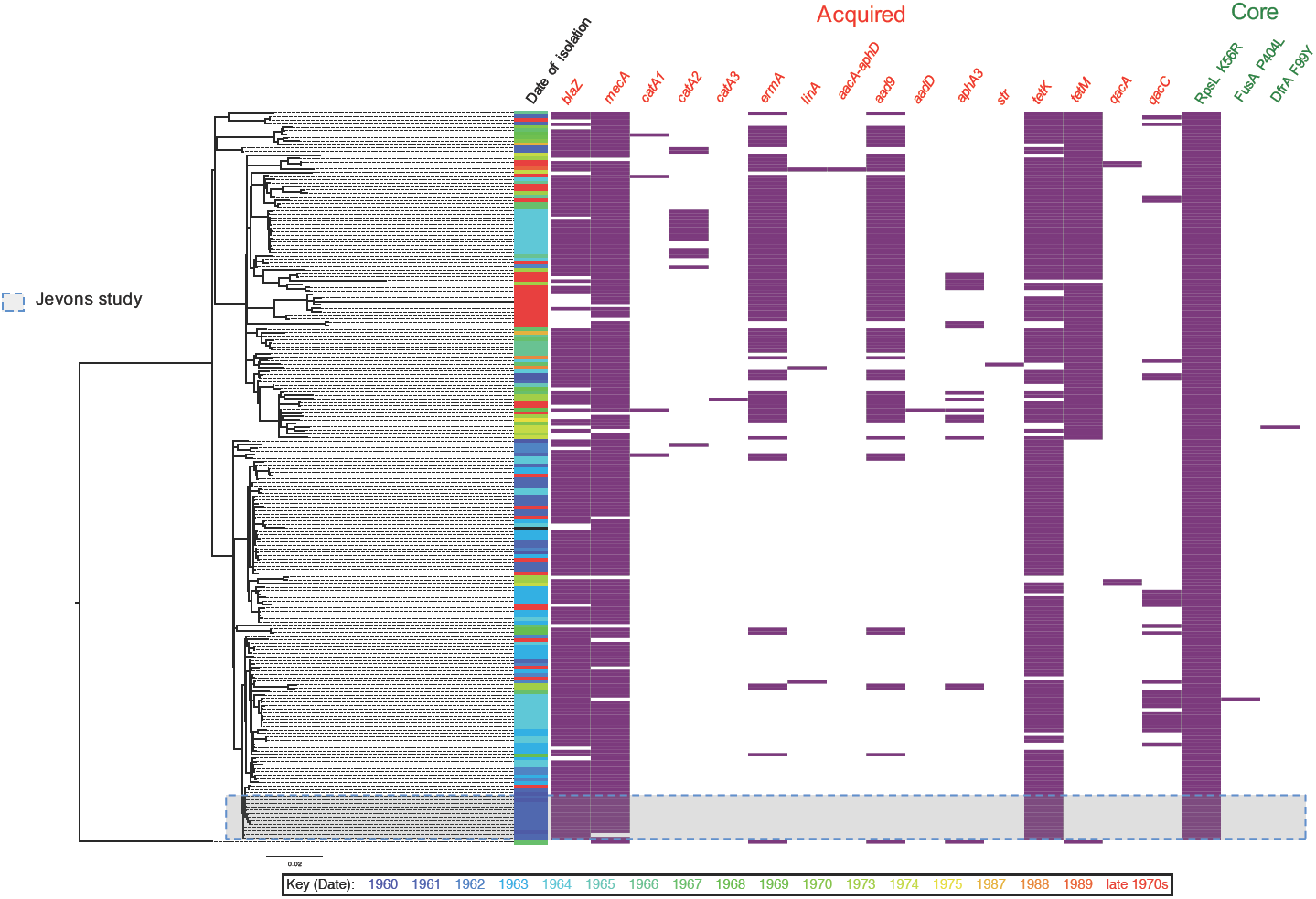
Distribution antibiotic resistance determinants in the archetypal MRSA clone. A maximum likelihood tree of historic MRSA isolates (n=209) plus the COL reference is displayed on the left, and the panels on the right indicate dates of isolation (coloured according to year, see key below for years), and the presence (purple boxes) and absence (white space) of genetic determinants responsible for antibiotic resistance in the genomes of the isolates. The identity of genetic determinants are listed at the top, and divided into acquired genes (red text), and core mutations (green text). The antibiotics linked to the genetic determinants for the acquired genes are: β-lactams *blaZ* and *mecA*; chloramphenicol, *catA1, catA2*, and *catA3*; erythromycin, *ermA*; clindamycin, *linA*; aminoglycosides, *aacA*-*aphD, aad9, aadD, aph3*A, and *str*; tetracycline, *tetM* and *tetK*; disinfectants, *qacA* and *qacC*; and for the core gene mutations: streptomycin substitution of arginine for a lysine at residue 56 (K56R) of the ribosomal protein *rpsL*; fusidic acid, substitution of a proline for a leucine at residue 406 (P404L) of the transcription elongation factor *fusA*; trimethoprim, substitution of an tyrosine for a phenylalanine at residue 99 (F99Y) of the dihydrofolate reductase *dfrA*.

### Evolution and emergence of methicillin resistance

To determine if the methicillin resistance emerged once or multiple times in the archetypal MRSA population we examined the variation within the SCC*mec* type I elements. In total, 194 variant sites were identified in 192 elements present in the collection. Analysis of the distribution of the variation within the elements suggested that some could be attributed to homologous recombination. Two regions contained the majority of the variation: 124 SNP sites were identified in the gene encoding the LPxTG surface protein *pls*, and 31 SNP sites within a 549 bp intergenic region between a hypothetical protein (SACOL0030) and a glycerophosphoryl diester phosphodiesterase (SACOL0031). Excluding these predicted recombination regions, 39 core variants sites across 28.6 kb distinguished the 192 elements, with half of the isolates (n=96) carrying an identical element. The maximum SNP distance distinguishing any two elements was eight SNPs, and phylogenetic analysis revealed that the elements present in the historic MRSA clone were closely related (Supplementary Figure 1) and shared a common evolutionary origin.

Our analysis of the evolutionary events surrounding the emergence of methicillin resistance in the archetypal MRSA lineage focused on a subset of 122 isolates that had precise dates of isolation (Supplementary Table 1). This enabled us to generate a robust Bayesian phylogeny and temporal calibration. Examining the distribution of the type I SCC*mec* variants (Figure 3A) within the context of a core genome phylogeny generated with BEAST (Figure 3B) reveal congruence between the phylogenetic relationships of the two. All of the canonical SNPs associated with the SCC*mec* genotypes could be singularly mapped onto nodes of the core phylogeny, suggesting that the variation observed in the SCC*mec* elements had occurred during expansion of the ST250 and ST247 populations. On the basis of this, we propose that a type I SCC*mec* element was acquired once in a single primordial development of methicillin resistance (Figure 3B), that could be dated back to the emergence of this clone.

We utilized Bayesian phylogenetic methods applying combinations of molecular clock models. The combination of an exponential population and relaxed-log normal clock model was found to be the best fit to our data based on Bayes factors using the harmonic mean estimator. This indicated the time to most recent common ancestor (TMRCA) of the ST250/ST247 population was 1946 (95% highest posterior density (HPD) 1938-1952) (Supplementary Figure 2), and therefore the time of acquisition of SCC*mec* was likely to around, or before this date. Notably, the time to most recent common ancestor of the type I SCC*mec* elements in these isolates based on a linear regression of a core SNP phylogeny was predicted to be early 1941 (Supplementary Figure 3).

To ensure that the Bayesian result was not an artifact of the clock or population models used in the analysis, we calculated the TMRCA for a range of model combinations and found that our chosen model exhibited a predicted TMRCA that was encompassed by the 95% HPDs of all other model combinations (Figure 4).

**Figure 3.**
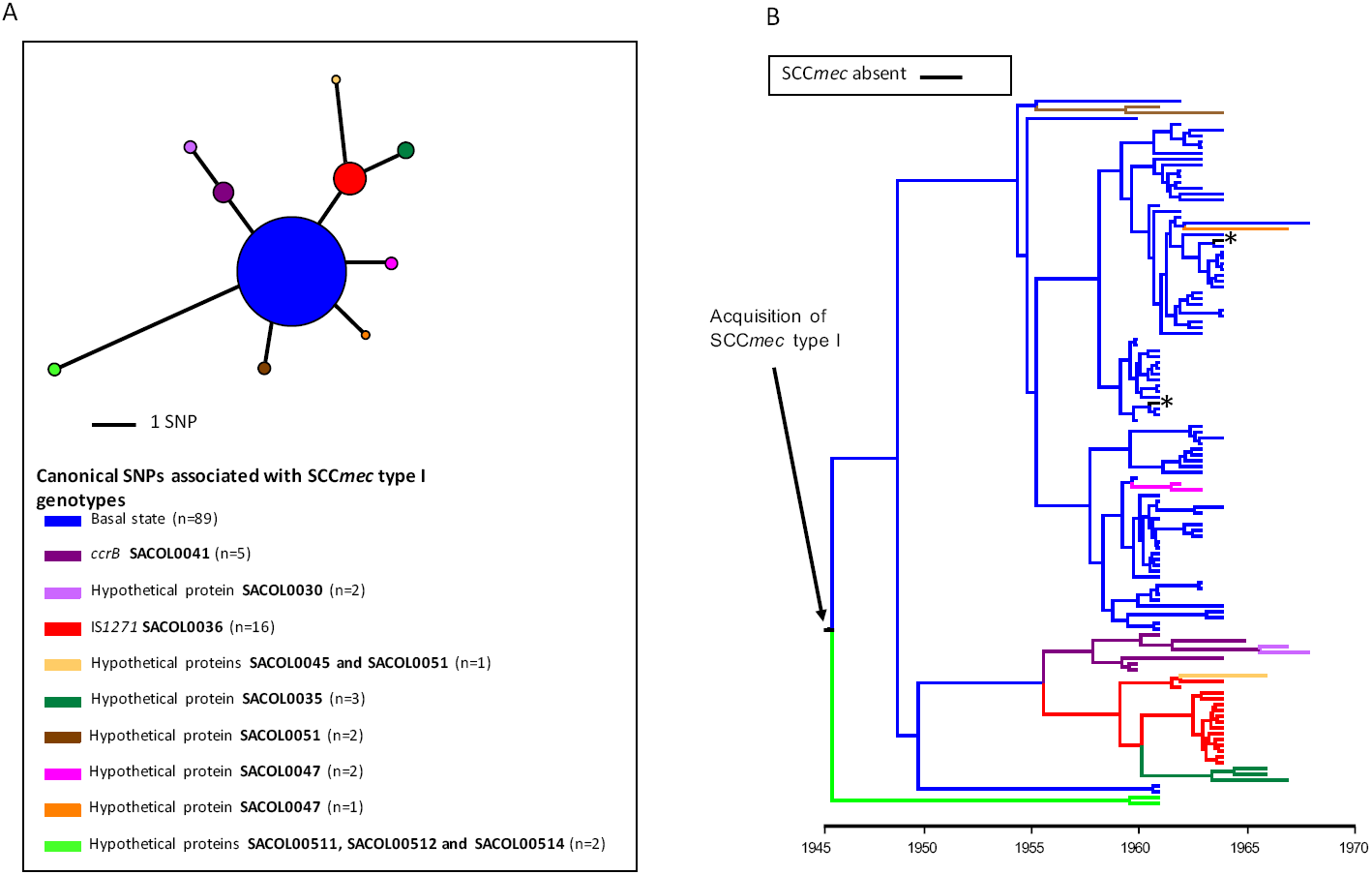
Diversity and distribution of SCC*mec* type I elements in the archetypal MRSA population. A. Parsimonious minimal spanning tree of SCC*mec* type I elements present in the archetypal MRSA isolates present in the clade credibility tree in panel B. The tree is built with core SNPs identified in the SCC*mec* type I elements, and excludes SNP in the *pls* gene that were predicted to have arisen by recombination. In total, 10 genotypes were observed, and the genetic events that distinguish each genotype from the founder genotype are indicated. The tree is centered on the majority genotype inferred as the founder population, and colour-coded according to their genotype. Black stars indicate isolates that lack the type I SCC*me*c element. The sizes of the circles illustrate the relative sizes of the genotype populations. The key below the tree describes the canonical SNPs differentiating SCC*mec* type I genotypes and the number of variants with that genotype. B. Maximum clade credibility tree of the archetypal MRSA clone population based on BEAST analysis. Tips of the tree are constrained by isolation dates; the time scale is shown below the tree. The tree is built with core genome SNPs from a subset of the total collection’s isolates (n=122), which had robust provenance and precise dates of isolation. The branches of the tree are coloured according to the genotype of the SCC*mec* type I element present in that strain (illustrated in panel 3A). Internal branches are coloured according to parsimonious reconstruction of the predicted genotype. Where terminal branches are black, this indicates the absence of an SCC*mec* element, which is predicted to reflect loss of the element. An arrow indicates the point in the phylogenetic reconstruction where an ancestral of type I SCC*mec* element was acquired. The root of the tree corresponds the basal node of the ST250/ST247 population in Figure 1 rooted by the RH12000692_7401696 outgroup. From the analysis the estimated mutation rate of population is 1.8×10^−6^ SNPs/site/year. This substitution rate falls within the reported ranges of multiple successful *S. aureus* lineages (Hsu et al. 2015) and therefore it is unlikely likely that long-term storage of the isolates has created any temporal artefacts.

**Figure 4.**
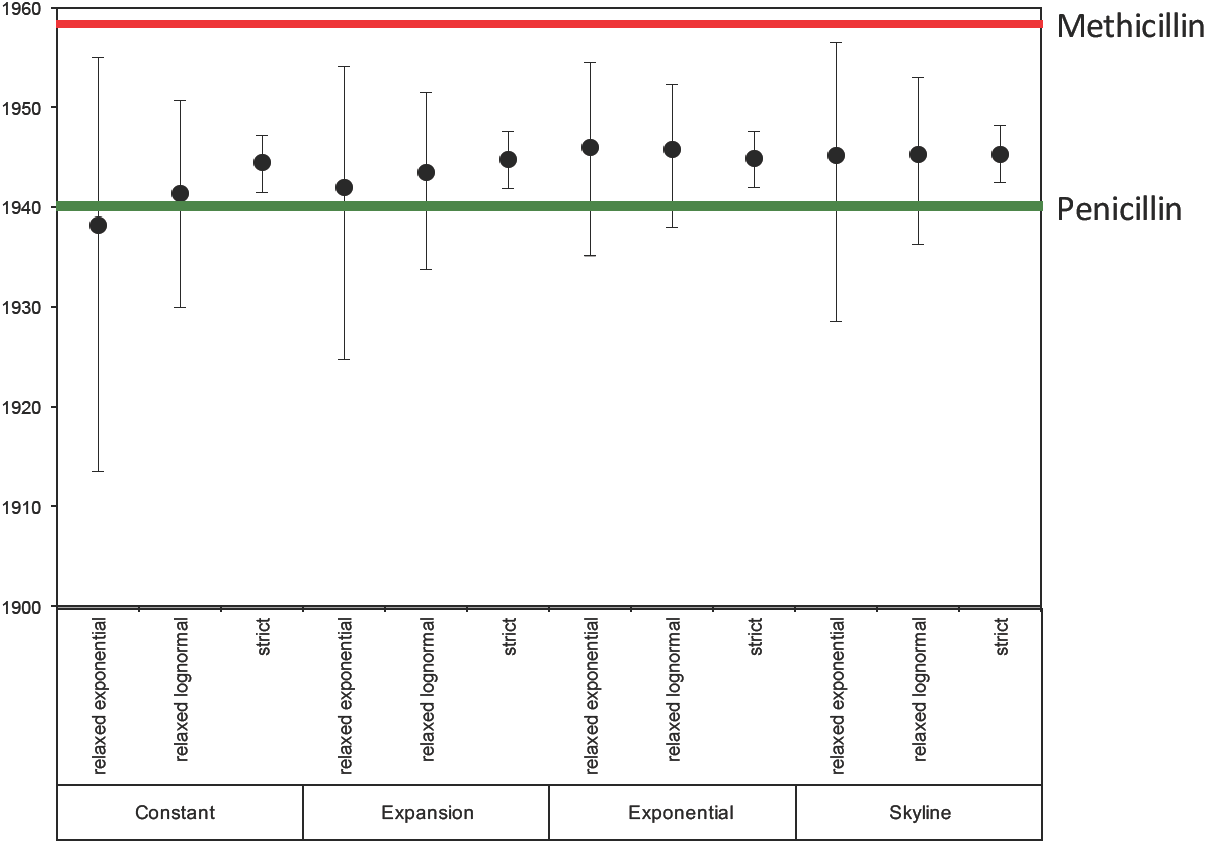
The time to most recent common ancestor (TMRCA) of the archetypal MRSA isolates under various combinations of clock and population model in BEAST. Plots showing mean (dots) TMRCA, and 95% HPD (highest posterior density) for the TMRCA are indicated. The dates of introduction of penicillin and methicillin into clinical use in the UK are indicate with green and red lines respectively.

## Discussion

This historic collection provides unique insights into of the evolution of the first MRSA lineage. Preserved for decades in their original freeze-dried state this large collection of strains representing the earliest MRSA clone has allowed us to reconstruct the evolutionary events leading to the emergence of MRSA. Using whole genome sequencing we have been able to shed light on the time when SCC*mec* first entered into *S. aureus*, and also to estimate how many times this is likely to have happened in the archaic MRSA population.

The origins of SCC*mec* almost certainly lie in coagulase negative staphylococci (CoNS) (Couto et al. 1996). *S. aureus* belonging to the ST250 background appear to have been the first recipient in the transfer from CoNS, but whether the element entered the ST250 population on multiple occasions, or as a single isolated event with subsequent propagation through the population has never been definitively resolved. A single entry of *mecA* followed by its evolution within the recipient background has been suggested (Kreiswirth et al. 1993). In order to clarify this we examined the variation present within the SCC*mec* elements in isolates throughout the population. The variation seen within SCC*mec* is predominantly in the *pls* gene which has been described before (Shore et al. 2005). Functionality of this 230 kDa cell wall-anchored (CWA) protein remains unclear, but its expression has been shown to reduce adhesion to host proteins as well as decrease invasiveness (Juuti et al. 2004). This LPXTG surface protein has a highly repetitive D/S rich structure making it a target for homologous recombination. As noted in other lineages the CWA proteins are subject to diversifying selection and exhibit diversity between and within *S. aureus* lineages (Santos-Júnior et al. 2016; Murphy et al. 2011). Removal of this variation reveals that the evolutionary history of the SCC*mec* elements was congruent with that of the strains carrying them, which points towards a single acquisition, rather than multiple or recurrent horizontal transmissions. Supporting this hypothesis is the observation of a mutation in *ccrB1* gene of the SCC*mec* type I element. The recombinase genes are required for both integration and excision from the chromosome. Specifically, CcrB is required for excision and the mutation present within this NCTCT10442 type I SCC*mec* element is believed to produce a non-functioning recombinase (Misiura et al. 2013; Noto and Archer 2006). Given that all the isolates in this collection have this frameshift mutation, this strongly supports the conclusions of the phylogenetic analysis, namely that a type I SCC*mec* was acquired once in the ST250 background, and then became fixed in the population due to defective recombinase apparatus that precluded excision.

One of the questions we sought to address in this study was what were the temporal events surrounding the emergence of MRSA. With the first reports of MRSA occurring only after introduction of methicillin in the UK in 1959 and Denmark in 1964 it seemed reasonable to conclude that resistance arose after the first clinical use of the drug, and resistance therefore developed in *S. aureus* as an adaptive response following exposure to the antibiotic. However the results presented in this communication are not consistent this conclusion, since the gene bestowing methicillin resistance was likely to have been acquired in the mid 1940s. It was during this period that β-lactamase mediated penicillin resistance was becoming widespread among clinical isolates of *S. aureus*. Within four years of the introduction of penicillin for the treatment of staphylococcal infections, the first penicillin-resistant *S. aureus* were being described in 1944 (Kirby 1944). In the years that followed the frequency of resistance in clinical isolates climbed steadily, such that by the time methicillin was introduced into clinical practice in 1960, resistance rates of 80% were common (Barber and Rozwadowska-Dowzenko 1948; Bauer et al. 1960).

Whilst the main genetic determinant associated with penicillin resistance in *S. aureus* is *blaZ, mecA* also encodes penicillin resistance via a different mechanism involving an alternative penicillin-binding protein, PBP2a (Hartman and Tomasz 1984; Ubukata et al. 1989). In the sequenced collection *blaZ* is widely distributed, albeit at a lower frequency than *mecA* (85.2% of isolates carry the *blaZ* gene in comparison to 95.2% for *mecA*) suggesting a selective advantage to possessing two distinct β-lactam resistance mechanisms. Based on the temporal calibration of the acquisition of *mecA*, it appears likely that methicillin resistance in *S. aureus* evolved long before this new β-lactam antibiotic was introduced. Thus it was the widespread use of penicillin, rather than methicillin, that was the driver for the emergence in the archaic MRSA clone.

Beyond β-lactams our analysis uncovered evidence for the strong selective impact that a number of different antibiotics have had on the evolution of the archaic MRSA clone. Several of the antibiotics, such as tetracycline, are prescribed in far lower amounts today in human medicine, than in the 1950s and 1960s, and resistance to these antibiotics in contemporary *S. aureus* from humans is relatively rare, which contrast the archaic MRSA population, where the distribution of tetracycline resistance determinants was widespread (Figure 2; 96% of isolates contained *tetK* or/and *tetM*) (Aanensen et al. 2016). In a prescient study examining the antibiotic consumption and rates of resistance in a hospital in the US in the 1950s, Bauer *et al*. provided evidence of a correlation between the two, where increasing usage of tetracycline was associated with an increase in the frequency of tetracycline resistance in isolates from inpatients (Bauer et al. 1960).

In addition to methicillin and tetracycline resistances, a key phenotypic marker of the archaic MRSA clone was non-susceptibility to streptomycin. In our analysis we identified a mutation predicted to confer streptomycin resistance occurring on the same branch of the tree in which we mapped the acquisition of the SCC*mec* element. This finding suggests that methicillin and streptomycin resistance both emerged in the archetypal MRSA progenitor population around the same time. Discovered in the early 1940s, streptomycin was demonstrated to have activity against Gram-positive pathogens, and was used in the UK in 1947 during the first ever randomized clinical trials studying the efficacy of streptomycin in the treatment of pulmonary TB (Schatz et al. 1944; Anonymous 1948). It therefore appears that the first MRSA clone emerged, and developed resistance to two of the earliest antibiotics - streptomycin and penicillin - almost immediately after the *S. aureus* population would have been first exposed to them.

At the time of its discovery, the incidence of MRSA in the general population is likely to have been very low. This is demonstrated by the fact that screening of over 5000 samples at Public Health England yielded only three methicillin resistant isolates. Therefore it is likely that when methicillin was introduced to circumvent penicillin resistance in *S. aureus*, it did not select for emergence of MRSA at that time, but instead provided the selective pressure, which drove the nosocomial spread of a pre-existing variant, at a time when infection control measures in UK hospitals were limited.

This study highlights the unintended consequences of widespread antibiotic use, and how when new drugs are introduced to bypass known resistance mechanisms, they may be already be rendered ineffective due to unrecognised adaptations accrued in response to prior selective pressures exerted by other antibiotics. This remains one of the many challenges in tackling the growing problem of AMR and serves to emphasise the importance of continual surveillance of pathogen populations for evidence of emerging adaptations and resistance patterns in the context of prescribing practice.

## Methods

### Bacterial isolates

Two hundred and nine isolates derived from the culture collections of *Staphylococcus aureus* reference laboratory, Public Health England, and isolates originating from the Statens Serum Institute collected and analysed by Profs Tomasz, Westh and de Lencastre. These correspond a collection of MRSA and MSSA isolates collected between 1960 and late 1980s in the UK and Denmark.

One hundred and eighty eight isolates preserved as freeze-dried cultures in the Health Protection England (HPA) Staphylococcal Reference Laboratory were resurrected and grown on solid media. Prior to the start of this study the Reference Laboratory sequence typed the all isolates from 1960 and 1961 using standard MLST techniques (Enright et al. 2000), and identified that the isolates belonged to clonal complex (CC) and were either sequence type (ST) 250 or 247.

Twenty one CC8 MRSA isolated in Denmark between 1964 and 1989 were also included in this study. These isolates originating from the Statens Serum Institute and had been previous sequence typed using standard MLST techniques (Enright et al. 2000). All isolates in this study were subsequently sequence typed from their whole genome sequence (WGS) data (see below).

### Genomic library preparation and sequencing

Genomic DNA was isolated using the Qiagen QIAcube system according to manufacturer’s protocol.

We prepared sequencing libraries from 500ng of DNA extracted from each MRSA isolate as previously described, with amplification using Kapa Hifi polymerase (Kapa Biosystems, Woburn, MA, USA) (Hsu et al. 2015). Tagged DNA libraries were created using a method adapted from a standard Illumina Indexing protocol, as described previously (Hsu et al. 2015). Whole genome sequencing was performed on the Illumina HiSeq 2000 platform with 100 bp paired-end reads. The Illumina sequence data has been submitted to the European Nucleotide Archive (ENA) and the accession numbers are provided in Supplementary Table 1.

### Bioinformatic and phylogenetic analysis

The sequence reads for each representative isolate (n=209) were aligned against the reference genome of the MRSA *S. aureus* COL (accession number: CP000046) (Gill et al. 2005) using SMALT (http://www.sanger.ac.uk/science/tools/smalt-0) and SNPs (single nucleotide polymorphisms) and indels (insertions/ deletions) identified as described previously (Hsu et al. 2015). Mobile genetic elements (MGEs) identified in the COL reference sequence by manual curation of BLASTN pairwise visualized in ACT (Carver et al. 2005). Regions of recombination within core genome and SCC*mec* element alignments were identified using Gubbins (http://github.com/sanger-pathogens/Gubbins) (Croucher et al. 2014). Phylogenetic reconstruction using core SNPs was performed with RAxML (Stamatakis 2014). Regions of high-SNP density corresponding to putative regions of recombination and those SNPs associated with horizontal gene transfer were excluded.

In order to investigate if the genomic data contained evidence of a temporal signal we used root to tip linear regression using Path-O-Gen v.1.4 (http://tree.bio.ed.ac.uk/software/tempest/)(Supplementary Figure 4). A core alignment for 122 isolates for which precise dates of isolation were available was used. MGEs and regions of predicted recombination along with homplastic SNPs within these isolates were then also excluded. To estimate evolutionary rates and time to most common recent ancestor (TMRCA) Bayesian phylogenetic reconstruction was performed using BEAST (v.1.7.4) (Drummond et al. 2012). A GTR model with a gamma correction for among-site rate variation was used, and all combinations of strict, relaxed lognormal, and relaxed exponential clock models and constant, exponential, expansion, and skyline population models were evaluated. For each, three independent chains were run for 100 million generations, sampling every 10 generations. On completion each model was checked for convergence, both by checking effective sample size (ESS) values were greater than 200 for key parameters, and by checking independent runs had converged on similar results. Models were compared for their fit to the data using Bayes factors based on the harmonic mean estimator as calculated by the program Tracer v.1.4 from the BEAST package. A burn-in of 10 million states was removed from each of the three independent runs of this model before combining the results from those runs with the logcombiner program from the BEAST package.

A previously described database of sequences of known resistance determinant genes, both horizontally acquired and core, was utilized as a resistome database (Holden et al. 2013). Fastq files from the 209 isolates were mapped to the resistome database using SRST2 (Inouye et al. 2014). SNPs in chromosomally encoded genes previously identified as being associated with antimicrobial resistance were then manually inspected to confirm the variation.

Multilocus Sequence Type (MLST) of isolates was predicted using SRST2 (Inouye et al. 2014).

### Data access

Short reads for all sequenced isolates have been submitted to the European Nucleotide Archive (ENA; http://www.ebi.ac.uk/ena/) under study accession number ERP001103. Individual accession numbers of sequences and assemblies for all isolates are listed in Supplementary Table 1.

## Acknowledgements

We thank Nick Thomson for reading the manuscript and useful discussions. We also thank the core sequencing and informatics teams at the Sanger Institute for their assistance and The Wellcome Trust for its support of the Sanger Institute Pathogen Genomics and Biology groups. S.D.B, J.P and M.T.G.H were supported by Wellcome Trust grant 098051. CPH was supported by Wellcome Trust Grant number 104241/z/14/z. Bioinformatics and Computational Biology analyses were supported by the University of St Andrews Bioinformatics Unit that is funded by a Wellcome Trust ISSF award (grant 097831/Z/11/Z). AK, MD and BP received funding from Public Health England.

## Author contributions

M.T.G.H. and C.P.H carried out data analyses, interpreted the data, and wrote the paper. B.P. and M.D. carried out data analyses and cultured the isolates and helped write the paper. S.D.B jointly conceived the project with A.M.K., and with A.T. and H.d.L˙. J.P. facilitated sequencing of the isolates, and helped write the paper. A.S.W. H.W., A.T., H.d.L., S.D.B and A.M.K. provided isolates and helped write the paper.

---

**Supplementary Figure 1**

**Maximum likelihood tree of SCC*mec* type I elements in historic MRSA isolates.**

The tree was constructed using variation in 38 SNP core sites identified in 192 isolates. The coloured branch labels indicate the isolates used in the parsimonious minimal spanning tree of SCC*mec* type I elements illustrated in Figure 3A and the temporal analysis illustrated in Figure 3B (also included in Supplementary Figures 1, 2 and 3), and are colour coded according to the genotypes displayed in this figure. The isolates not included in the temporal analysis are in indicated in black text.

**Supplementary Figure 2**

**Posterior support of maximum clade credibility trees of the historic MRSA population based on BEAST analysis (as illustrated in Figure 3B).**

Internal branches are colored according to their posterior support (see figure for key).

**Supplementary Figure 3**

**Linear regression of the root-to-tip distances of historic MRSA SCC*mec* type I elements.**

The isolates used (n=122) are those indicated in the Supplementary Figure 4 and used for the BEAST analysis (Figure 3B). The analysis was carried out using Path-O-Gen v1.4 (http://tree.bio.ed.ac.uk/software/pathogen/) with a best-fit root from the maximum likelihood tree and the dates of isolation. The plot contains straight-line best fit of the root-to-tip divergence for each of the isolates, with a correlation coefficient of 0.5408 and an R^2^ of 0.2925. The time to most recent common ancestor for the SCC*mec* type I elements in the archaic clone isolates examined is 1941.

**Supplementary Figure 4**

**Linear regression of the root-to-tip distances of the archetypal MRSA clone population used for BEAST analysis.**

The analysis was carried out using Path-O-Gen v1.4 (http://tree.bio.ed.ac.uk/software/pathogen/) with a best-fit root from the maximum likelihood tree and the dates of isolation. The plot contains straight-line best fit of the root-to-tip divergence for each of the isolates, with a correlation coefficient of 0.7907 and an R^2^ of 0.6525. The time to most recent common ancestor for the whole population was estimated to be 1947, consistent with the Bayesian analyses (as illustrated in Figure 3B).

**Supplementary Table 1**

**Isolate metadata, including dates and origin isolates where available, mapping and assembly statistics, and genotype information.**

**Supplementary Table 2**

**Isolates from the original description of MRSA.**

Minimum inhibitory concentration (MIC) to celbenin (methicillin) derived from the original description by P. Jevons, published in the BMJ in 1961. MIC values represent the variation noted between colonies. Expected range of sensitivity to celbenin in coagulase positive staphylococci 1.25 - 2.5μg/ml. Acquired antibiotic resistance genes and core resistance mutations identified in the genomes are indicated.

